# Enterovirus-A71 exploits Rab11 to recruit chaperones for virus morphogenesis

**DOI:** 10.1101/2023.08.05.552141

**Authors:** Qing Yong Ng, Vikneswari Mahendran, Ze Qin Lim, Justin Jang Hann Chu, Vincent Tk Chow, Newman Siu Kwan Sze, Sylvie Alonso

## Abstract

Enterovirus 71 (EV-A71) causes Hand, Foot and Mouth Disease (HFMD) in children and has been associated with neurological complications. A siRNA screen in EV-A71 infected-motor neurons identified small GTPase Rab11a as a pro-viral host factor. Rab11a and Rab11b isoforms were interchangeably exploited by strains from major EV-A71 genogroups and Coxsackievirus 16, another major causative agent of HFMD. We showed that Rab11 did not play a role in viral entry, IRES-mediated protein translation, or viral genome replication, although it co-localized with replication organelles. GTPase-defective Rab11 mutants had no effect on EV-A71 replication, ruling out Rab11 involvement in intracellular trafficking of viral or host components. Instead, reduced VP2:VP0 ratio in siRab11-treated cells supported a role in provirion maturation. Co-immunoprecipitation and mass spectrometry revealed chaperones as Rab11 top interacting partners during infection. Among which, CCT8 subunit of the chaperone complex TRiC/CCT was found to interact with viral structural proteins specifically. Together, this study describes a novel, unconventional role for Rab11 during viral infection, where it participates in the complex process of virus morphogenesis by recruiting essential chaperone proteins.

## INTRODUCTION

After the near complete eradication of its close cousin Poliovirus, Enterovirus-A71 (EV-A71) has emerged as a public health concern, particularly among paediatric patients [1, 2]. EV-A71 belongs to the *Picornaviridae* family and, together with Coxsackievirus A16 (CVA16), is one of the main causative agents of Hand Foot Mouth Disease (HFMD). HFMD is highly infectious and typically affects children aged 5 and below, but also the elderly and immunocompromised individuals. Millions of HFMD cases are reported every year worldwide, with recurring outbreaks in the Asia Pacific region every 2 to 3 years [1]. Uncomplicated, self-limiting HFMD manifests as sore throat, fever, skin rash, ulcers in the mouth and blisters on the soles and palms. More rarely, HFMD may result in polio-like neurological symptoms, such as acute flaccid paralysis and encephalomyelitis that can be fatal or lead to long-term cognitive and motor deficits [3-5]. EV-A71 infections have been more frequently associated with such devastating complications granting this highly transmissible virus the status of one of the most medically concerning neurotropic virus.

EV-A71 is a non-enveloped virus that harbours a 7.4kb single-stranded RNA genome within its capsid. The viral RNA encodes for a single polyprotein that is proteolytically cleaved into four viral structural proteins (VP1-VP4) and seven non-structural proteins (2A-2C, 3A-3D) [6]. EV-A71 strains are grouped under 11 sub-genotypes namely A, B1-B5, C1-C4, largely characterised by their highly variable VP1 sequences. New sub-genotypes, such as D and E, have been introduced over the years, illustrating the fast-evolving nature of EV-A71 genomes, fuelled by high mutation rate and recombination events among co-circulating strains [7, 8].

There is currently no effective therapeutic drug against HFMD [5]. Treatment is supportive in nature and often combined with public health measures such as school and playground closure to try and limit transmission [9]. Those measures force parents to take leave to look after their children, adding an economic dimension to this disease in affected societies. Three EV-A71 vaccines have been approved in China, and consist of formalin-inactivated C4 virus formulations, the predominant sub-genotype in China [10, 11]. The effectiveness of these monovalent vaccines however has yet to be evaluated outside China, in regions where the predominant circulating sub-genotype is not C4, and/or where several EV-A71 sub-genotypes co-circulate.

The lack of effective treatment options against HFMD in general and EV-A71 in particular, is mainly attributed to limited research efforts to understand the molecular mechanisms of pathogenesis. Specifically, identification of the viral determinants and host factors involved in EV-A71 fitness and virulence are key to the rationale design of effective interventions. Using a proteomics approach, our group has previously reported two novel host factors, namely prohibitin-1 (PHB) and peripherin (PRPH) that are exploited by EV-A71 during its infection cycle in neuronal cells. While PHB was found to be closely associated with viral replication complexes located at the mitochondrial membrane [12], intermediate neurofilament PRPH was shown to support virus entry and viral genome replication through interactions with structural and non-structural viral components [13]. The latter study also led to identify small GTP-binding protein Rac1, a druggable host factor.

Pursuing our efforts to identify host factors involved in EV-A71 infection cycle, we have undertaken a siRNA screen that targets 112 host genes encoding for proteins involved in intracellular membrane trafficking. Indeed, membrane trafficking pathways are commonly targeted by viruses, as they need to transport their viral components to various subcellular compartments for genome replication, protein translation, post-translational modifications, virus assembly and virus exit [14, 15]. EV-A71 has been known to induce massive membrane re-arrangements to support its replication, and a number of host proteins have been associated with these events [15]. Here, we have identified 16 and 5 genes that encode for pro- and anti-viral factors, respectively. Among the pro-viral candidates, knockdown of Rab11a expression resulted in the most significantly reduced viral titers, implying that this host factor plays an important role during EV-A71 infection cycle.

A member of the Rab family, Rab11a is a small GTPase involved in the exocytic and late endosomal recycling pathways. Similar to other Rabs, Rab11 is activated when a guanine exchange factor (GEF) replaces the bound GDP with GTP. GTP-bound Rab11 interacts with its protein partners known as Rab11-family interacting proteins (FIPs), which subsequently allows Rab11 to engage with a series of other proteins that help Rab11-bound vesicles to move towards their destined subcellular location [16, 17]. Upon GTP hydrolysis, Rab11 then interacts with yet another group of proteins which promote activities such as membrane fusion [17]. Three isoforms of Rab11 have been reported, namely Rab11a, Rab11b and Rab25. Rab11a and Rab11b isoforms share 90% nucleotide sequence homology, while similarity with Rab25 is around 60%. All three isoforms are known to be involved in the recycling pathways, but they are also believed to have distinct cellular functions [18]. They are also differentially expressed whereby Rab11a is ubiquitously expressed, while Rab11b expression is limited to the brain, testis and heart, and Rab25 is restricted to the epithelial cells in lungs, colon and kidney [18].

A number of viruses have been reported to exploit Rab11 during infection, with influenza virus being the most well studied [19-24]. Rab11 has been described to promote aggregation and transport at the plasma membrane of the eight viral RNA segments during the viral assembly step [19, 24]. In contrast, little is known about the involvement of Rab11 during EV-A71 infection. Current literature suggests that Rab11 is exploited by enteroviruses for modulating the cholesterol level in replication organelles (ROs) to create a microenvironment favourable for viral RNA synthesis [25]. Such cholesterol re-routing is associated to PI4KB, which is recruited to the ROs by viral protein 3A through GBF1 or ACBD3 proteins [25-29]. In this scenario, Rab11 is believed to facilitate the transport of extracellular/plasma membrane cholesterol towards the ROs.

Here, we have combined various experimental approaches to dissect the role of Rab11 during EV-A71 infection, including genetics, biochemistry, dynamic confocal imaging and mass spectrometry techniques. Our data suggest that the main role of Rab11 during EV-A71 infection is to support the maturation and assembly of newly formed virus particles, rather than facilitating trafficking activities.

## MATERIAL AND METHODS

### Cell lines and virus strains

Human rhabdomyosarcoma (RD) cell line (American Type Culture Collection (ATCC), CCL-136), mouse motor neuron-like hybrid cell line NSC-34 (CELLutions Biosystems, CLU140), and human neuroblastoma SH-SY5Y cell line (ATCC #CRL-2266) were maintained in Dulbecco’s Modified Eagle’s Medium (DMEM) supplemented with 10% fetal bovine serum (FBS) (GIBCO) at 37°C with 5% CO2. Enterovirus-A71 (EV-A71) clinical isolates of sub-genotype B4 (strain S41, Genbank accession number AF316321), B5 (EVGP-18-254 strain, Genbank accession number OQ571388), C1 (EVGP-18-331 strain, Genbank accession number OQ571387), C2 (C2 strain, Genbank accession number NUH0075/SIN/08), and CVA16 (CA16-G-10, Accession number U05876), were plaque purified and propagated in RD cells maintained in DMEM supplemented with 2% FBS. Harvested culture supernatants containing the virus particles were aliquoted and stored at −80°C.

### siRNA knock-down (Reverse transfection)

Appropriate dilutions of both siRNAs (Table S1) and transfection reagents (Dharmafect-1 for knockdown in RD and NSC34 cells, and Dharmafect-4 for knockdown in SH-SY5Y cells) were performed in MEM-RS (Cytiva, Cat #SH30564.01). Equal volume of diluted siRNAs and transfection reagents were mixed and incubated for 30mins at room temperature (RT). After 30mins, 20 uL or 100uL of siRNA-Dharmafect mixture were added to each 24-well or 96-well respectively. 10^5^ cells in 400uL or 2.5 x 10^4^ cells in 80uL of culture medium were then added to each 24-well and 96-well respectively. The plate was incubated for 48h at 37°C, 5% CO2 before further analysis or treatments were performed.

### siRNA library screening

Mouse Membrane Trafficking siRNA library (Dharmacon, Cat # 115505) was reconstituted using DPEC-treated water (Invitrogen, Cat # AM9906). For each 96-well (Thermo Scientific, Cat #243656), 2.5 x 10^4^ NSC34 cells were reversed transfected with 50nM of each siRNA pool from the library. At 48h post transfection, the cells were infected with EV-A71 sub-genotype B4 Strain S41 at a multiplicity of infection (MOI) of 30, and the culture supernatants were collected at 48 hours post-infection (h.p.i). The viral titers were determined by plaque assay and expressed as percentage of viral titer reduction compared to siRNA non-template control (NTC). Two independent screening campaigns were performed.

### Virus Kinetics Assay

Reverse siRNA-transfected cells were infected with S41 virus at MOI 0.1 (SH-SY5Y and RD cells), or MOI 30 (NSC34 cells). After two washes with 1X PBS, fresh 1X DMEM supplemented with 2% FBS was added to the cells, and the plates were further incubated at 37°C, 5% CO_2_. At the indicated time points, the infected cells and their culture supernatants were harvested for Western blot analysis and plaque assay respectively. The culture supernatants were centrifuged at 4,000*g* for 10mins to remove any cells debris before the virus titers were determined by plaque assay. Concurrently, the cell monolayers were washed once with 1X PBS, and harvested by gentle scrapping. The three replicate wells were pooled together and centrifuged at 2,500*g* for 10mins. The cells pellets were resuspended in M-PER reagent (Thermo Scientific, Cat #78503) containing protease inhibitor (Thermo scientific, Cat #87786) and EDTA. After incubation on ice for 20mins, the lysates were centrifuged at 14,000*g* for 15mins, and the clarified lysates were stored at −20°C for further analysis by Western blot.

### Cell viability assay

Cell viability was determined by AlamarBlue assay. Briefly, each well was washed once with 1X PBS prior to addition of AlamarBlue™ reagent (Invitrogen; Cat # DAL1025; diluted 1:10 in complete growth medium). The plates were then incubated at 37°C, 5% CO_2_ for 1hr. Fluorescence signals were measured using Tecan SPARK plate reader (Ex570nm and Em585nm). The percentage of cell viability was determined in reference to untreated control.

### Western Blot

Total protein content in cell lysates was determined by Bradford assay. 10ug of total protein per sample were mixed with reducing Laemmli buffer (Bio-Rad, Cat # 161074) and heated at 95°C for 10mins. SDS gel electrophoresis was then run at 90V for 2hrs. For total protein content, precast stain-free gel (Bio-Rad, Cat #4568045) was imaged using Bio-Rad ChemiDoc. The proteins were next transferred onto nitrocellulose membrane (Bio-Rad, Cat #170427) using Bio-Rad Trans-Blot Turbo System. The membranes were then blocked in blocking buffer (5% blocking-grade milk in TBS-T) for 1hr, followed by overnight incubation with respective primary antibodies diluted in blocking buffer (Table S2). The membranes were washed thrice with TBS-T before incubation with the appropriate secondary antibodies diluted at 1:3,000 in blocking buffer for 1hr (Table S2). The membranes were washed thrice in TBS-T before addition of chemiluminescent substrate (Thermo Scientific, Cat # 34076) and imaging using X-ray or Bio-Rad ChemiDoc. Images were analysed using ImageJ or ImageLab for semi-quantification.

### Plaque assay

RD cells were seeded in 24-well plates (10^5^ cells per well) and incubated overnight at 37°C, 5% CO2. Clarified virus-containing culture supernatants were subjected to 10-fold serial dilution in 2% FBS-DMEM, and 100uL of each dilution (10^-1^ to 10^-6^) were added per well. Three technical replicates were performed for each dilution. The cells were incubated for 1hr at 37°C, 5% CO_2_. The cells were then washed twice with 1X PBS before overlaying with DMEM containing 1% carboxymethyl cellulose (Sigma Aldrich, Cat # 419303) and 2% FBS. The plates were incubated at 37°C, 5% CO_2_ for 72hrs before removing the overlay medium to fix and stain the cells using 0.02% crystal violet containing 4% PFA for 1h. Plaques were scored visually at the appropriate dilution and viral titers were expressed as plaque forming units per mL (PFU/mL).

### Entry by-pass assay

EV-A71 S41 RNA was extracted from 400uL of infected culture supernatant using viral extraction kit (Qiagen, Cat # 52906). SH-SY5Y cells were transfected with siRab11 mix, targeting both Rab11a and Rab11b isoforms. At 48h post-transfection (h.p.t), 500ng of purified viral RNA and 2uL of lipofectamine 2000 (Invitrogen, Cat #11668019) were each diluted in 50uL Opti-MEM (Gibco, Cat # 51985034), mixed together and incubated at RT for 30mins before being added to the siRNA-treated SH-SY5Y cells. The cells were incubated for another 24hrs at 37°C, 5% CO_2_ before the supernatants were harvested for viral titer determination by plaque assay.

### LucEV-A71 replicon and Bicistronic construct Assays

*E. coli* strains harbouring LucEV-A71 replicon or bicistronic construct [30, 31] were cultured in LB broth containing 50ug/mL kanamycin (Gibco, Cat # 11815032). The plasmids were extracted using QIAprep Spin Miniprep Kit (QIAGEN, Cat # 27104) according to the manufacturer’s instructions. LucEV-A71 assay: The plasmid was linearised using *Mlu*1 restriction enzyme (NEB, Cat # R3198S), and purified using Phenol:Chloroform:Isoamyl Alcohol (PCI 24:25:1; Sigma-Aldrich, Cat # 77617). In-vitro Transcription was performed using 1ug of linearised plasmid with MEGAscript® T7 Transcription Kit (Thermo Scientific, Cat # AM1334) according to the manufacturer’s protocol. The suspension was cleaned up using Phenol:Chloroform:Isoamyl (CPI 24:25:1) and chloroform (Sigma, Cat # C2432). SH-SY5Y cells, reverse-transfected with siRab11 mix or siNTC, were incubated at 37°C, 5% CO_2_ for 48hrs. After 48h, 500ng of LucEV-A71 transcript and 2uL of lipofectamine 2000 (Invitrogen, Cat #11668019) were each diluted in 50uL Opti-MEM (Gibco, Cat # 51985034) and mixed together. The resulting 100uL of RNA-lipofectamine mixture was incubated at RT for 30mins before adding it to the siRNA-treated SH-SY5Y cells. The plate was incubated for another 24hrs at 37°C, 5% CO_2_ before measurement of luciferase activity using Nano-Glo® Luciferase Assay (Promega, Cat # N1110) and Tecan Spark plate reader.

Bicistronic assay: siRab11-treated cells were incubated at 37°C, 5% CO_2_ for 48hrs. After 48h, 500ng of plasmid and 2uL of lipofectamine 2000 (Invitrogen, Cat #11668019) were each diluted in 50uL Opti-MEM (Gibco, Cat # 51985034) and mixed together. The resulting 100uL of DNA-lipofectamine mixture was incubated at RT for 30mins before adding it to the siRNA-treated SH-SY5Y cells. The plates were incubated for another 24hrs at 37°C, 5% CO_2_ before measurement of Renilla and Firefly luciferase activities using Dual-Glo® Luciferase Assay System (Promega, Cat #E2920) and Tecan SPARK plate reader. Results were expressed as FLuc:RLuc ratio.

### Immunofluorescence Assay (IFA) and Proximity Ligation Assay (PLA)

SH-SY5Y cells (10^5^) were seeded onto coverslips placed into a 24-well plate and were incubated at 37°C, 5% CO_2_ overnight. Where indicated, cells were first transfected with 500ng of relevant plasmids (eGFP-Rab11a DN/CA/WT) using Lipofectamine™ 3000 (Invitrogen; Cat #L3000015) according to the manufacturer’s instructions. The cells were then incubated at 37°C, 5% CO_2_ for 24hr, before infection with S41 virus at MOI 0.1. At 48 h.p.i, the cells were fixed with 4% PFA for 15mins at RT, permeabilized using 0.1% Tween-20/PBS for 15mins and washed thrice with 1X PBS.

IFA: The coverslips were blocked in 2% BSA in PBS for 1hr at 37°C, followed by incubation for 1hr at 37°C with the relevant primary antibodies (Table S3), which were diluted in blocking buffer. The coverslips were washed thrice with 1X PBS, followed by incubation with appropriate secondary antibodies (Table S3) for 1hr at 37°C. For compartment staining, coverslips were incubated with the respective compartment markers for 1hr at 37°C after washing thrice in 1X PBS. The coverslips were washed thrice again in 1X PBS, followed by incubation with Hoechst 33342 (Invitrogen, Cat # R37605) for 15mins at RT. The coverslips were then mounted onto microscope slides using DABCO (Sigma Aldrich, Cat # 10981-100ML) before being imaged either with Olympus IX81 fluorescence microscope or Olympus FV3000 Confocal microscope.

PLA: PLA was carried out using the Duolink™ In Situ Red Starter Kit Mouse/Rabbit (Sigma-Aldrich, Cat #DUO92101-1KT). Briefly, the coverslips containing fixed cells (as described above) were blocked using blocking buffer, followed by incubation with the appropriate mouse and rabbit antibodies (Table 3) at 37°C for 1 hour. The coverslips were then washed thrice with Buffer A, before incubation with the two PLA probes for 1 hour at 37°C. After 3 washes with Buffer A, Ligase was added and incubated for 30mins. The ligase was washed off with buffer A before incubation with polymerase for 100 mins. The coverslips were then mounted onto microscope slides using Duolink™ In Situ Mounting Medium with DAPI, and the slides were imaged with Olympus IX81 or Olympus FV3000 Confocal microscope.

### Co-immunoprecipitation (Co-IP)

EV-A71 infected and uninfected SH-SY5Y cells were scrapped off T175 tissue culture flasks, and were centrifuged at 2,500*g* for 10mins. The cell pellets were washed once with 1X PBS before being lysed in MPER reagent containing protease inhibitors (Thermo Scientific; Cat # 87785) and 5mM EDTA. The lysates were then spun down at 14,000*g* for 15mins. Dynabeads crossed-linked to the appropriate antibodies were added to the clarified lysates and incubated at 4°C for 3hrs. The beads were washed thrice in 1X PBS and subjected to elution using 1% SDS. The eluted fractions were then mixed with Laemmli buffer and heated at 95°C for 10mins before being analysed by Western blot.

### Co-IP/Mass Spectrometry

Co-IP was performed as described above, with the elution performed by heating the beads in Laemmli buffer at 70°C for 10mins. The eluted samples were then subjected to 8-20% gradient SDS-PAGE. The protein bands were then excised and subjected to in-gel trypsin digestion. The digested peptides were separated and analysed using Dionex Ultimate 3000 RSLCnano system coupled to a Q

Exactive instrument (Thermo Fisher Scientific, MA, USA). Separation was performed on a Dionex EASY-Spray 75 μm × 10 cm column packed with PepMap C18 3 μm, 100 Å (Thermo Fisher Scientific) using solvent A (0.1% formic acid) and solvent B (0.1% formic acid in 100% ACN) at flow rate of 300 nL/min with a 60-min gradient. Peptides were then analyzed on a Q Exactive apparatus with an EASY nanospray source (Thermo Fisher Scientific) at an electrospray potential of 1.5 kV. Raw data files were processed and converted to mascot generic file (mgf) format using Proteome Discoverer 1.4 (Thermo Fisher Scientific). The mgf files were then used for protein sequence database search using Mascot algorithm to identify proteins. Further analysis of identified proteins were done by employing both PantherDB and StringDB platforms. The mass spectrometry proteomics data have been deposited to the ProteomeXchange Consortium via the PRIDE [32] partner repository with the dataset identifier PXD043870.

*Note to the Reviewers/Editor: The dataset will be made publicly available after the manuscript has been accepted. Meanwhile, the dataset can be accessed during the reviewing process via* http://www.ebi.ac.uk/pride/archive *with the following credentials:*

***Username:***

***Password:*** *8jYh4jw4*

### Over-expression of Rab11a DN/CA/WT in SH-SY5Y cells

eGFP-Rab11a dominant negative (DN) and constitutively active (CA) plasmids [24] were obtained from Madison University, USA. Plasmid harbouring eGFP-Rab11a-WT was obtained by mutagenesis of eGFP-Rab11a-DN construct with Phusion® High-Fidelity DNA Polymerase (Thermo Scientific, Cat # F530S) using primers as follows: Forward ggtgttggaaagaGtaatctcctgtct; Reverse agacaggagattaCtctttccaacacc, where the capital letter features the point mutation introduced). These plasmids were transformed into *E. coli*, which were cultured in LB broth supplemented with 50ug/mL of kanamycin. The plasmids were extracted using Qiagen HiSpeed Plasmid Maxi Kit. SH-SY5Y cells (1.5−10^7^) were seeded into a T175 flask and incubated at 37°C, 5% CO_2_ overnight. Each T175 flask was transfected with mixtures consisting of 52.5ug of plasmid in 131.25uL of Lipofectamine 3000 and 87.5uL of p3000 reagent. The flasks were incubated at 37°C, 5% CO_2_ for 24h. At 24 h.p.t, cells were trypsinized and subsequently sorted using BD FACSAria Fusion cell sorter. The GFP^+^ sorted cells were seeded into a 48-well plate (7.5−10^4^ cells per well). Cells were allowed to rest at 37°C, 5% CO_2_ for 24h. The cells were then transfected with 50nM siRab11b and were further incubated at 37°C, 5% CO_2_ for 48h before infection with S41 at MOI 0.1. At 18 h.p.i, the supernatants and cells were harvested for viral titer determination by plaque assay and Western blot anlaysis, respectively.

### Statistical analyses

All statistical analyses were performed using GraphPad Prism 9.0. Mann-Whitney statistical test was performed for comparing independent data pairs, while Kruskal-Wallis H test was employed for the comparison between 3 or more independent groups. The degree of significance was indicated by asterisks with *p<0.05, **p<0.01, ***p<0.001, ****p<0.0001, ns: not significant.

## RESULTS

### siRNA screening identifies Rab11a as a pro-viral factor during EV71 infection in motor neurons

Motor neuron-like NSC34 cells were subjected to a siRNA screen targeting 120 genes involved in membrane trafficking. We have previously described NSC34 cells as an *in vitro* infection model predictive of EV-A71 *in vivo* neurovirulence [33]. Viral titers in the culture supernatant were measured by plaque assay providing information on the overall effect of the siRNA-mediated knockdown (KD) on virus replication. Prohibitin-1 encoding gene (Phb) was used as positive control as we previously showed that PHB is involved in both entry and post-entry steps of EV-A71 infection cycle in NSC34 cells [12]. Non-targeting scramble siRNA control (NTC) was used as negative control. The screening was performed twice independently and a total of 21 hits were obtained that resulted in viral titer reduction greater than 50% compared to the siNTC-treated control (Fig. 1A). Among which, Rac1 was identified, which was previously reported by us as a pro-viral host factor during EV-A71 infection in neuronal cells [13], thereby validating our screen.

**Figure 1:**
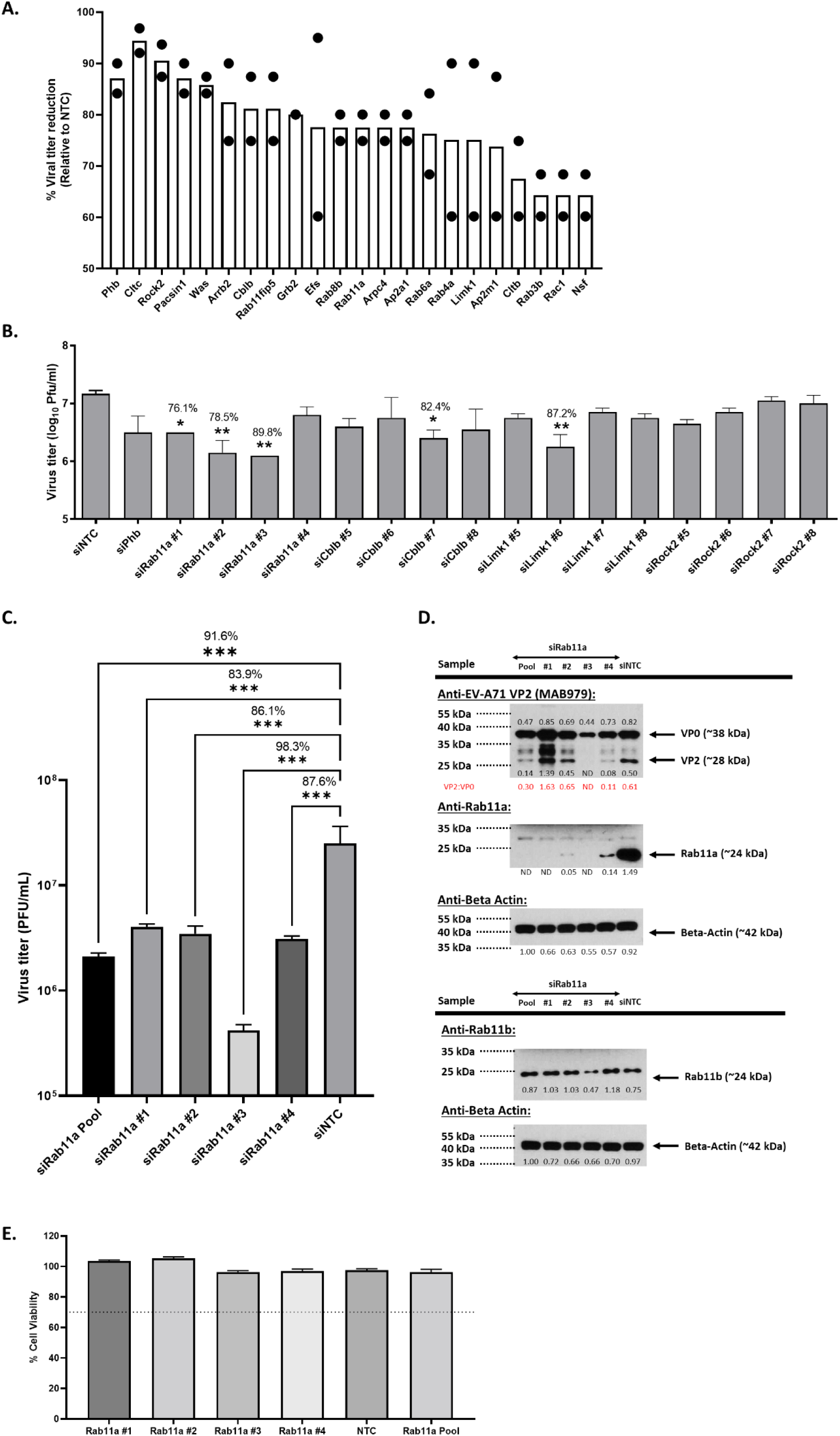
Identification of Rab11a as a pro-viral factor through screening of membrane trafficking siRNA library in NSC34 cells. **(A)** NSC34 cells were transfected with the siRNA library pools, then infected with EV-A71 S41 (MOI 30) 48 hours later. At 48 h.p.i, the culture supernatants were harvested to determine the viral titers by plaque assay. Results are expressed as the percentage of virus titer reduction obtained for each siRNA treatment compared to Non-targeting siRNA non-targeting control (NTC). Genes with virus titer reduction >50% in two independent screening experiments are shown. **(B)** NSC34 cells were transfected with deconvoluted siRNA targeting genes encoding Rab11a, Cblb, Limk1 and Rock 2, followed by infection with EV-A71 S41 (MOI 30) 48 hours later. At 48 h.p.i, virus titers in the culture supernatants were determined by plaque assay. **(C&D)** Further validation of Rab11a as a pro-viral factor using deconvoluted siRNAs. NSC34 cells were transfected with pool or deconvoluted siRNA targeting Rab11a, or with siNTC, followed by infection with EV-A71 S41 (MOI 30) at 48h.p.t. The culture supernatants and cell lysates were harvested at 48h.p.i. **(C)** Virus titers in the culture supernatants were determined by plaque assay. **(D)** The cell lysates were subjected to Western blot analysis using anti-VP2, anti-Rab11a, anti-Rab11b and anti B-actin antibodies. Band intensities were normalised to Beta-actin and relative to NTC. **(E)** AlamarBlue assay was performed on uninfected siRNA-treated cells at 48 hours post-transfection. The readout for each treatment was relative to non-siRNA-treated cells to determine the percentage of cell viability. Percentage above 70% (indicated by the dotted line) was considered non-cytotoxic. Statistical analysis was performed for **(B)** and **(C)** using Kruskal-Wallis test against siNTC treatment (*p<0.05, **p<0.01, ***p<0.001, ****p<0.0001). The percentages of virus titer reduction for siRNA treatments that were statistically different from siNTC are indicated above the asterisk.

Four genes encoding for Limk1, Rock2, Rab11a and Cblb were selected for downstream validation using deconvoluted siRNA, based on their known physiological role in cytoskeleton (Limk1 and Rock 2), recycling endosomal pathway (Rab11a) and proteasome degradation (Cblb). Rab11a KD resulted in significant viral titer reduction for three of the four siRNA species used (Fig. 1B). In contrast, KD of Cblb and Limk1 expression led to viral titer reduction for only one siRNA species, while no significant viral titer reduction was seen with Rock2 deconvoluted siRNAs. These data prompted us to select Rab11a for further characterization.

Deconvoluted siRab11a KD experiment was repeated and confirmed significant reduction in viral titers compared to siNTC, thus validating that Rab11a is a pro-viral host factor during EV-A71 infection in NSC34 cells (Fig. 1C). In addition, Western blot analysis of the cell lysates confirmed that Rab11a expression was effectively knocked down, which correlated with reduced VP2 signals except for siRNA#1 (Fig. 1D). Interestingly, treatment with siRNA#3 was found to also impact the expression of Rab11b isoform, which correlated with even greater reduction in viral titer (Fig. 1C) and VP2 signal (Fig. 1D), compared to the other siRNA-treated samples. This observation hence suggested that Rab11a and Rab11b isoforms may both be exploited by EV-A71 during its infection cycle in NSC34 cells. Interestingly, we also noticed that in Rab11 KD samples (except for siRNA#1), the ratio VP2:VP0 was clearly lower compared to NTC control (Fig. 1D), hence suggesting a potential role for Rab11a/b in VP0 maturation.

### Rab11a and Rab11b isoforms are exploited by EV-A71 and CVA16 during infection in human cell lines

To further explore the role of Rab11a during EV-A71 infection, a similar siRNA KD approach was performed in human rhabdomyosarcoma (RD) cell line, which has been widely used to study EV-A71 pathogenesis. Unlike NSC34 cells, Rab11a KD in RD cells using Rab11a-specific siRNA pool and individual siRNA species did not lead to significant viral titer reduction (Fig. 2A), except for siRNA#9 for which significant reduction in both viral titer and VP2 signal were observed (Fig. 2A&B). Interestingly, treatment with siRNA#9 led to reduced expression in both Rab11a and Rab11b isoforms, unlike the other siRNA species which impacted Rab11a expression only (Fig. 2B). This finding thus further supported the idea that both Rab11a and Rab11b are exploited interchangeably by EV-A71 during infection. To confirm this hypothesis, Rab11b-specific KD was performed in RD cells. Consistently, only treatment with siRNA #7, which resulted in significantly reduced expression of both Rab11a and Rab11b led to reduced viral titers in the culture supernatant, and reduced VP2 signal intensity in the cell lysates (Fig. 2C&D). To further demonstrate that co-KD expression of both Rab11a and Rab11b was responsible for reduced viral titers and VP2 intensity, RD cells were co-treated with siRab11a #10 and siRab11b #9 (A10B9 mix). While individual treatment with each siRNA species did not affect the viral titer and VP2 band intensity, co-treatment with both siRNA species led to significantly reduced viral titer and VP2 band intensity (Fig. 2C&D), hence demonstrating redundant functional role of Rab11a and Rab11b isoforms during EV-A71 infection. Furthermore, here again samples that were effectively knocked down for Rab11a and Rab11b, displayed lower VP2:VP0 ratio, hinting at a role for these proteins in VP0 maturation process. Similar observations were made in human neuroblastoma SH-SY5Y cells treated with siRab11a#10, siRba11b#9 or A10B9 mix (Supplemental Fig. S1).

**Figure 2.**
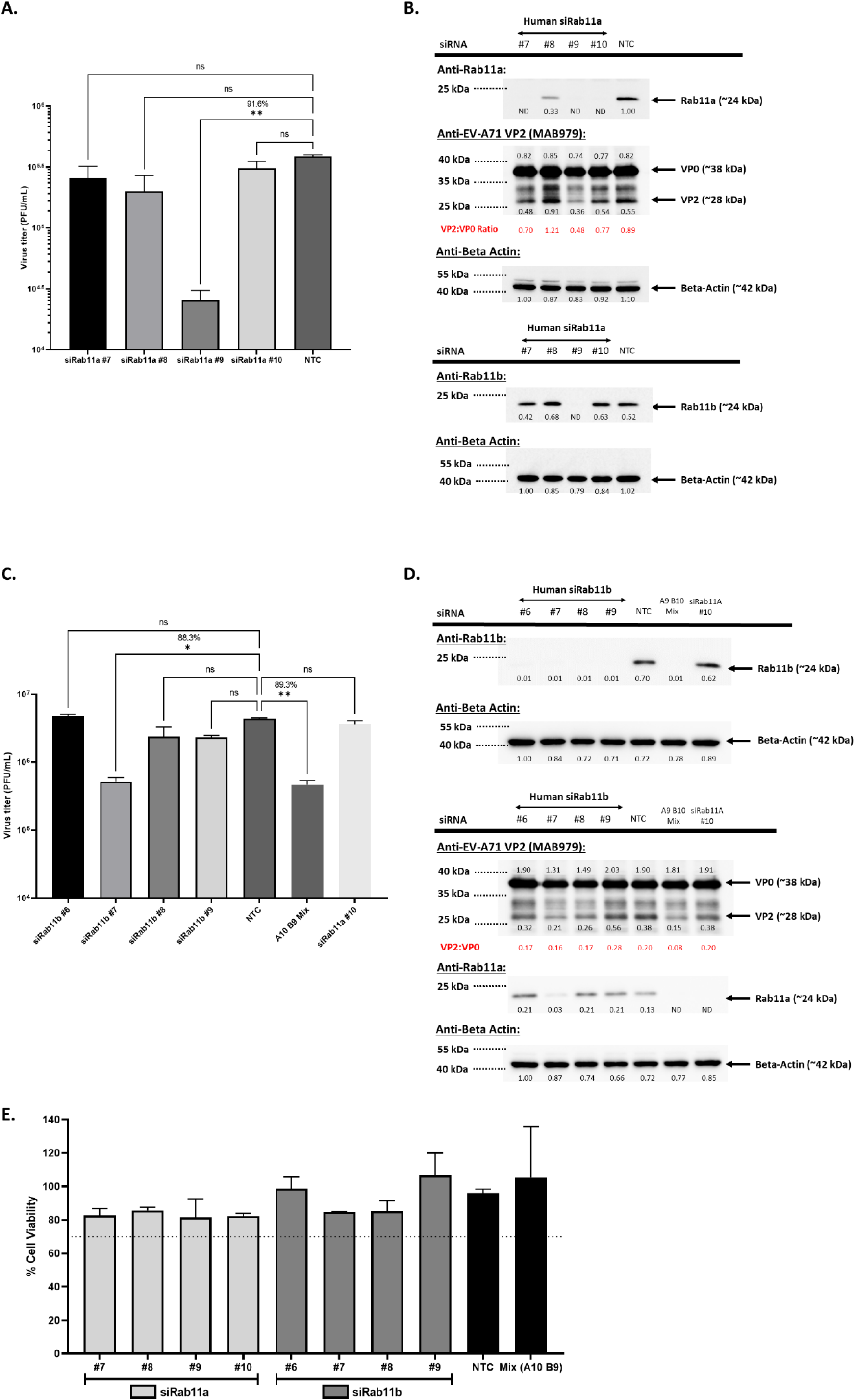
Effects of Rab11a and Rab11b KD on EV-A71 infection in RD cells. RD cells were transfected with deconvoluted human siRNA targeting Rab11a **(A&B)** or Rab11b **(C&D)**. At 48h.p.t, these cells were next infected with EV-A71 S41 (MOI 0.1). At 24 h.p.i, the culture supernatants and cell lysates were harvested. **(A&C)** Virus titers in the culture supernatants were determined by plaque assay. Statistical analysis was performed using Kruskal-Wallis test against siNTC treatment (*p<0.05, **p<0.01, ***p<0.001, ****p<0.0001). The percentages of virus titer reduction compared to NTC are indicated above the asterisk. **(B&D)** The cell lysates were subjected to Western blot analysis using anti-VP2, anti-Rab11a, anti-Rab11b and anti B-actin antibodies. Band intensities were normalised to Beta-actin and relative to NTC. **(E)** AlamarBlue assay was performed on uninfected siRNA-treated cells at 48 hours post-transfection. The readout for each treatment was relative to non siRNA-treated cells to determine the percentage of cell viability. Percentage above 70% (indicated by the dotted line) was considered non-cytotoxic.

We next investigated the importance of Rab11a/b during infection with EV-A71 strains representative of various EV-A71 sub-genogroups. SH-SY5Y cells were treated with siRab11a#10, siRab11b#9 or A10B9 mix before infection. Results indicated that significant reduction in viral titer and VP2 signal (and VP2:VP0 ratio) was observed in siA10B9-treated cells but not in cells treated with individual siRab11a#10 and siRab11b#9 (Fig. S2). The same observations were made with Coxsackievirus A16 (CVA16), a close cousin of EV-A71 and another main causative agent of HFMD (Fig. S2).

Taken together, the data supported that Rab11a and Rab11b isoforms are exploited by all major EV-A71 genogroups and CVA16 during infection, which suggests a conserved and important role for these proteins during infection. The consistent drop in VP2:VP0 ratio in Rab11-KD samples also suggested that Rab11 proteins may be involved in VP0 maturation.

### Rab11a/b is not involved in viral entry, genome replication and viral translation

To understand the role of Rab11a/b during EV-A71 (and CVA16) infection cycle, various assays were performed. Using an entry bypass assay, we first demonstrated that Rab11a/b was not involved in the entry step (including receptor binding, internalization and virus uncoating). In this assay purified viral RNA genome was directly transfected into siRab11-treated SH-SY5Y cells (resulting in KD of both Rab11a and Rab11b), thereby bypassing the entry step. Significant viral titer reduction was still observed (Fig. 3A), hence suggesting that Rab11a/b has minimal role in the viral entry step of EV-A71 infection cycle.

**Figure 3:**
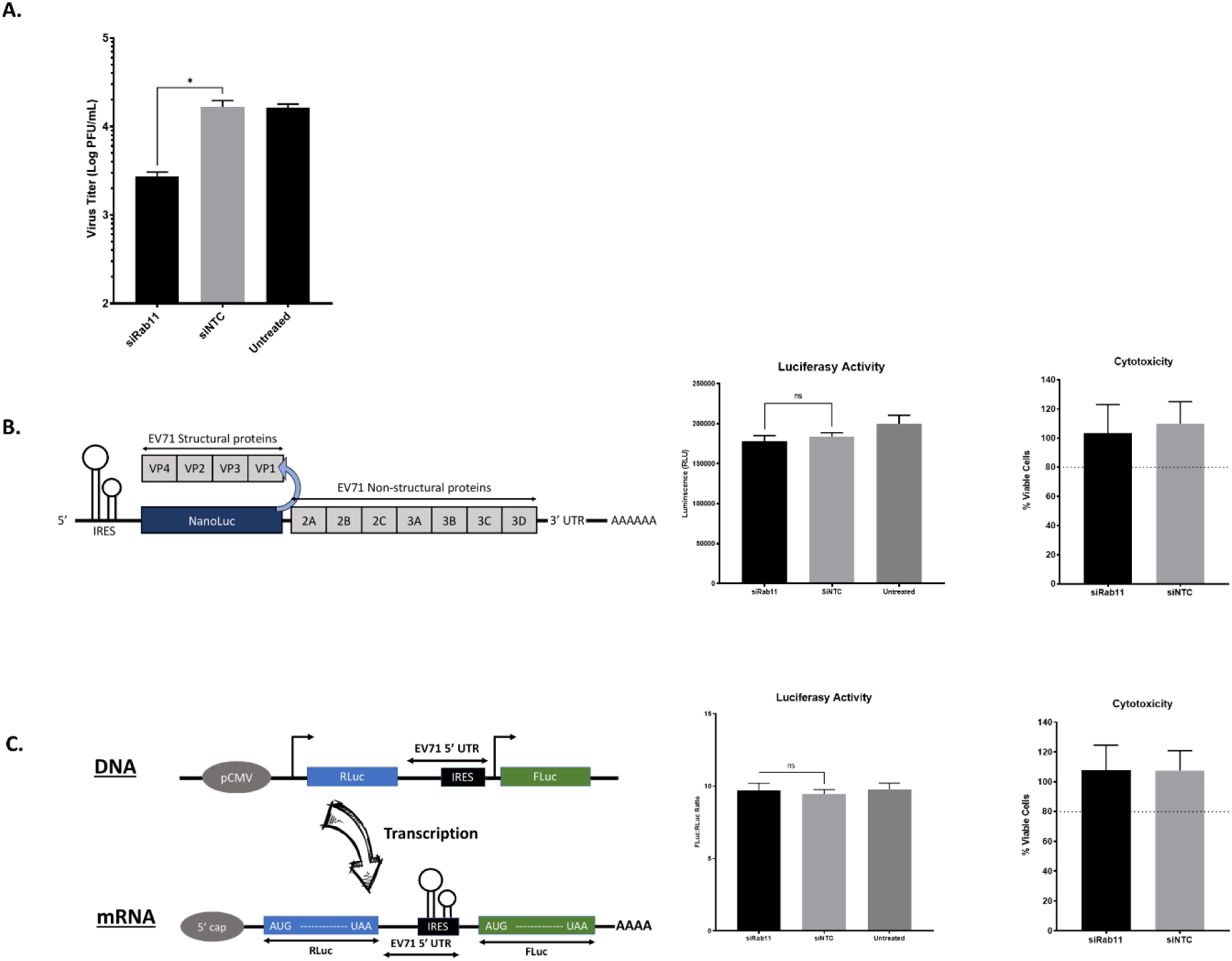
Role of Rab11 in viral entry, genome replication and viral protein translation. **(A)** Entry-bypass assay. SH-SY5Y cells were transfected with siRab11 or siNTC, followed by transfection with 500ng of EV-A71 S41 RNA genome. The culture supernatants were harvested 24h post-transfection, and the viral titers were determined by plaque assay. **(B)** EV71-Luc replicon assay. SH-SY5Y cells were treated with siRab11 or siNTC, then transfected with 500ng of *in vitro* transcribed EV71-Luc RNA. The luciferase activity and cytotoxicity were measured at 24 hours post-transfection. **(C)** EV71-Bicistronic reporter assay. SH-SY5Y cells were treated with siRab11 or siNTC, then transfected with 500ng of EV71 Bicistronic plasmid construct. Renilla and Firefly luciferase activities, and cytotoxicity (alamarBlue assay) were measured at 24 hours post-transfection. Statistical analysis was performed using Kruskal-Wallis test against siNTC treatment (*p<0.05, **p<0.01, ***p<0.001, ****p<0.0001).

We next assessed the role of Rab11a/b in viral RNA genome replication. A Luc-EVA71 replicon was employed where EV-A71 structural genes (P1 region) were replaced by NanoGlo luciferase encoding gene (Fig. 3B) [30]. siRab11-treated SH-SY5Y cells were transfected with *in vitro* transcribed Luc-EVA71 RNA, and the luciferase signal was compared to NTC-treated SH-SY5Y cells. No significant difference in signal intensity between siRab11-treated and NTC-treated cell cultures was observed (Fig. 3B), thus indicating that Rab11a/b is not involved in the initial round of viral RNA genome translation and replication by viral RNA-dependent RNA polymerase. We further confirmed that Rab11a/b does not play a role in viral genome translation by using a bi-cistronic construct, where a firefly luciferase-encoding gene is transcribed from a CMV promoter and contains EV-A71 cap-independent IRES for protein translation (Fig. 3C) [31]. No significant difference in luciferase activity was measured between siRab11-treated and NTC-treated cells (Fig. 3C), thus indicating that Rab11a/b does not contribute to IRES-dependent translation.

Together, the data supported that Rab11a/b is neither involved in viral entry step, nor does it contribute to viral genome replication and translation activities.

### Rab11a/b interacts with viral structural and non-structural proteins in various subcellular compartments

To gain further insights into the role of Rab11a/b during EV-A71 infection cycle, we sought to investigate physical interactions between Rab11a/b and viral proteins by performing Proximity Ligation assay (PLA). Results showed that both Rab11 isoforms co-localised with all the viral components tested including VP2, 3C, 3D, 3CD, and dsRNA (Fig. 4A&B). Furthermore, pull-down experiment using anti-Rab11a antibody showed that VP0, VP1, VP2, 3C, 3D, and 3CD were detected in the Rab11a pulldown samples (Fig. 4C), thus confirming close interactions between Rab11a protein and structural and non-structural viral proteins.

**Figure 4.**
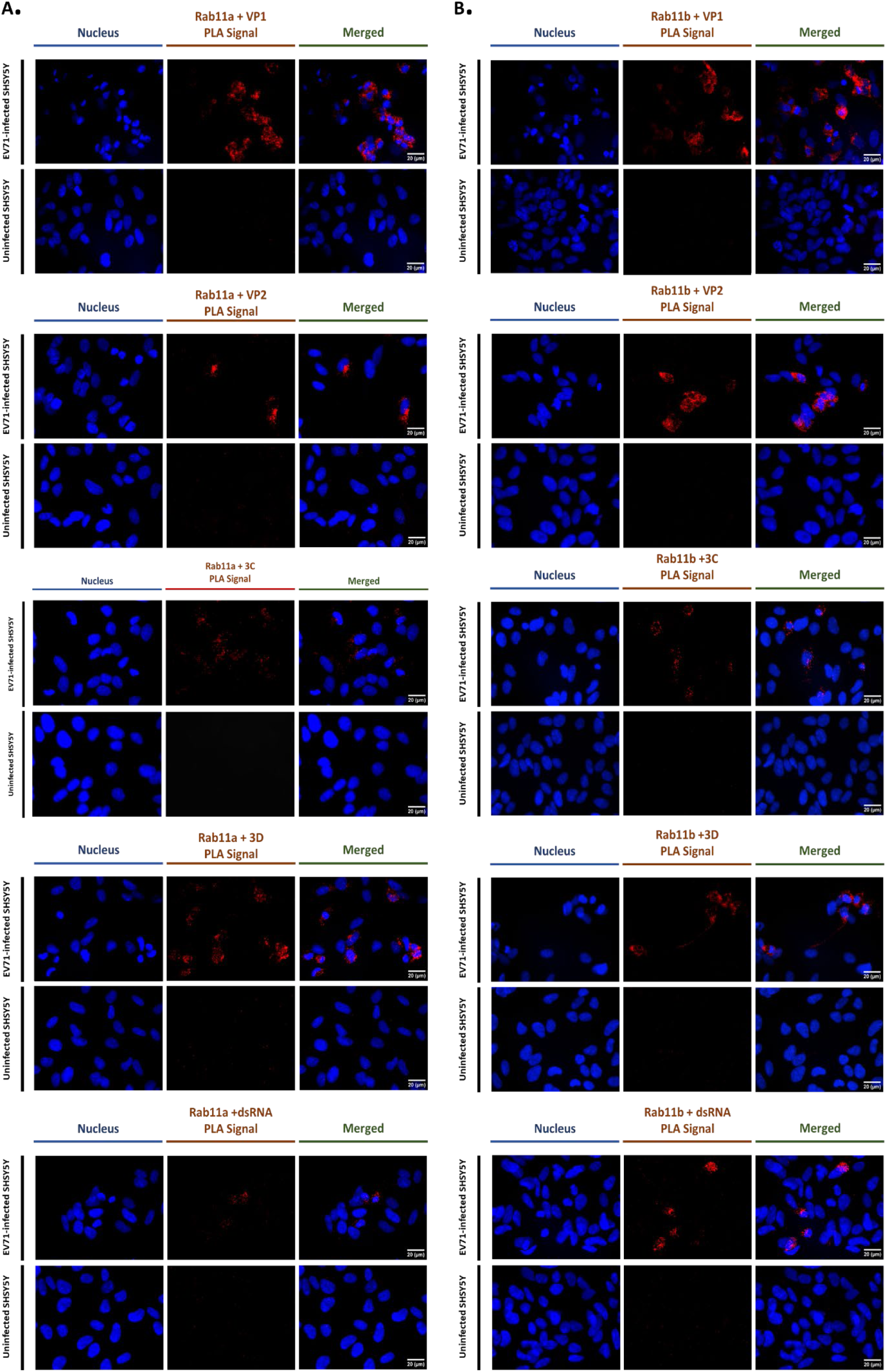
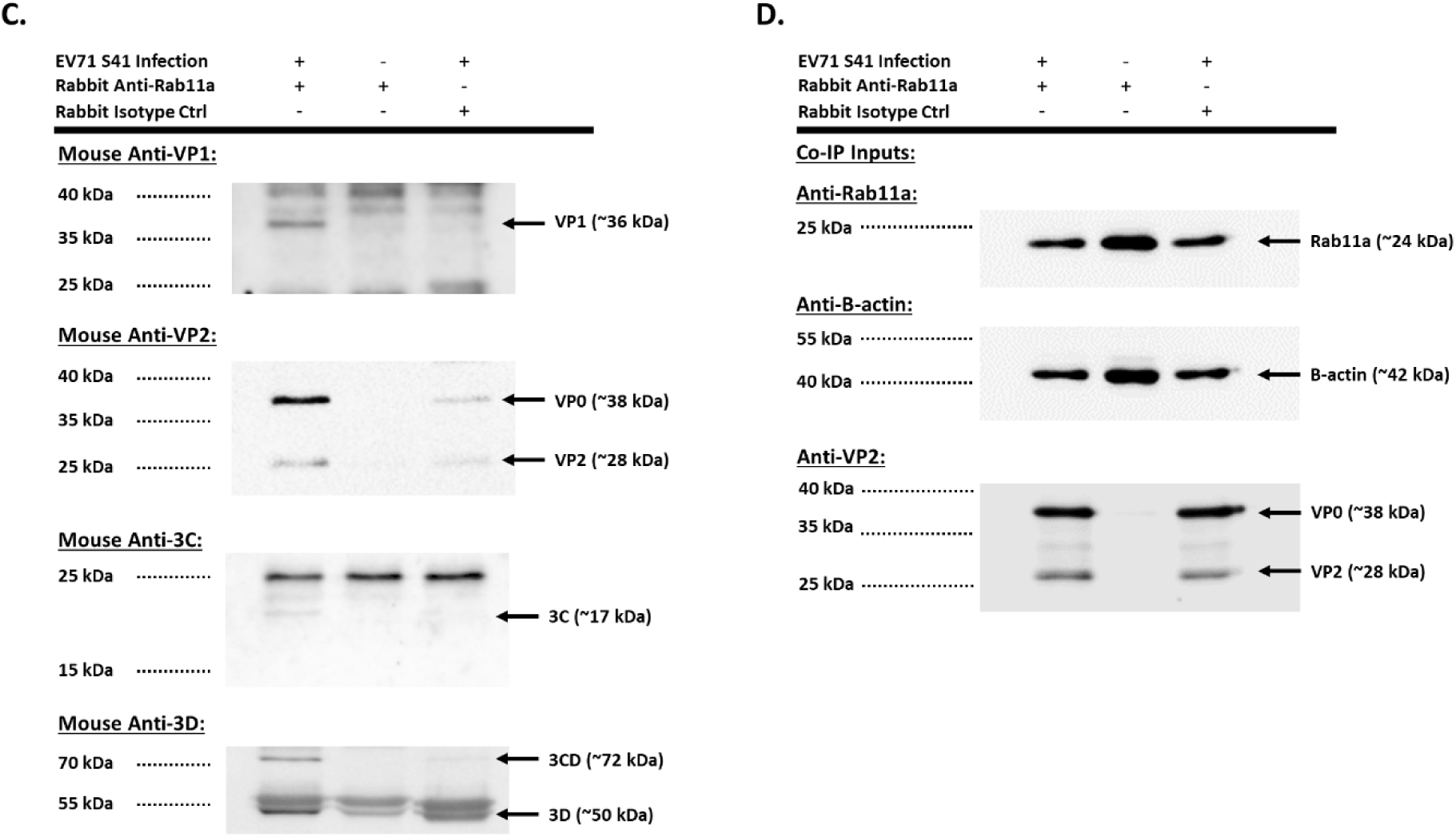
Co-localization and interaction of Rab11a and Rab11b with viral components. **(A&B)** Proximity ligation assay (PLA). SH-SY5Y cells were infected with EV-A71 S41 (MOI 0.1). At 24 h.p.i., the cells were fixed and permeabilized before staining with anti-Rab11a **(A)** or with anti-Rab11b **(B)**, and anti-VP2, anti-VP1, anti-3D, anti-3C or anti-dsRNA antibodies. The cells were further stained with secondary antibodies conjugated with DNA probes. Ligation and polymerase chain reaction were then carried out for signal amplification. Nuclei were stained with DAPI. Images were captured at 60X magnification under Olympus IX81 microscope. **(C&D)** Co-immunoprecipitation (Co-IP). SH-SY5Y cells were infected with EV-A71 S41 (MOI 0.1). At 24 h.p.i., the cells were lysed for immunoprecipitation using anti-Rab11a or rabbit IgG isotype control monoclonal antibodies. **(C)** The pulldown samples were subjected to Western blot analysis using anti-VP1, anti-VP2, anti-3C and anti-3D antibodies. **(D)** Infected and uninfected cell lysates (Co-IP inputs) were analysed by Western blot prior to Co-IP.

Since Rab11a/b are known to be involved in the exocytic and late endosomal recycling pathways, we hypothesized that these proteins may help transport the various viral components to their destined subcellular locations, for viral particle assembly and maturation. To address this hypothesis, multiplex confocal imaging was performed at various time points post-infection to track the dynamic interactions between Rab11a and viral proteins (VP2/VP0 and 3C/3CD) in various subcellular compartments. Results showed that interaction between Rab11a and VP2/VP0 could be detected from 6 hours post-infection onwards in the endoplasmic reticulum (ER), Golgi apparatus (GA), and small recycling endosomes (Fig. 5). Similar observations were made when probing interactions between Rab11a and 3C/3CD (Fig. S3). Furthermore, Pearson correlation coefficient (PCC) values were determined to quantify the strength of co-localization between Rab11a and the viral proteins of interest, whereby the higher the PCC values the stronger the co-localization between targets. Results indicated that co-localization between Rab11a and VP2/VP0 was moderate and constant across the various subcellular compartments and across time (Fig. 5D). Similar observations were made with Rab11a and 3C/3CD interactions (Fig. S3). Furthermore, we also evaluated interactions between Rab11a and VP1, VP2, 3D, 3C or dsRNA in the three subcellular compartments at 6 h.p.i. Again, comparable PCC values were obtained with all the viral components tested and across the three cellular compartments (Fig. S4).

**Figure 5.**
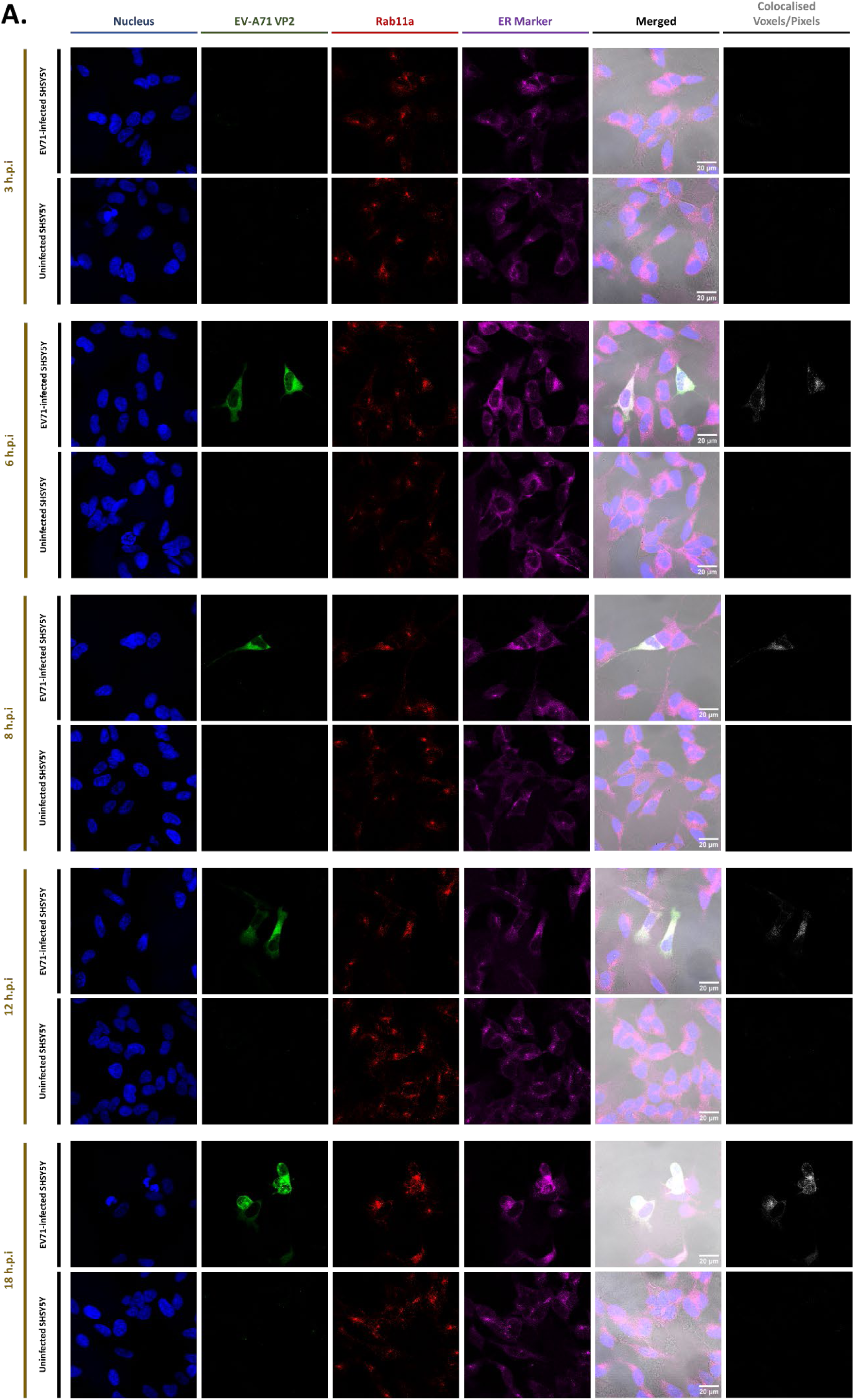
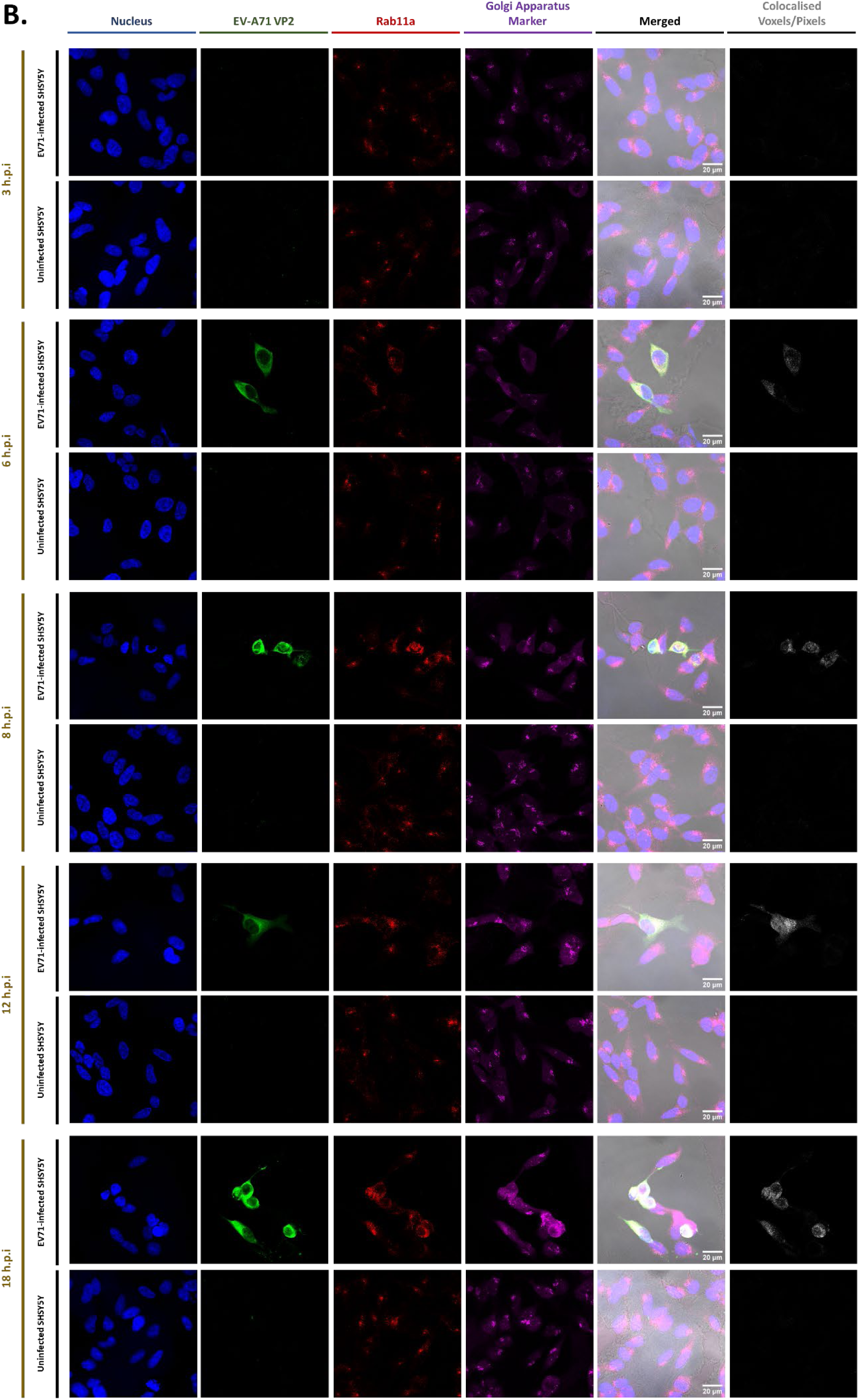
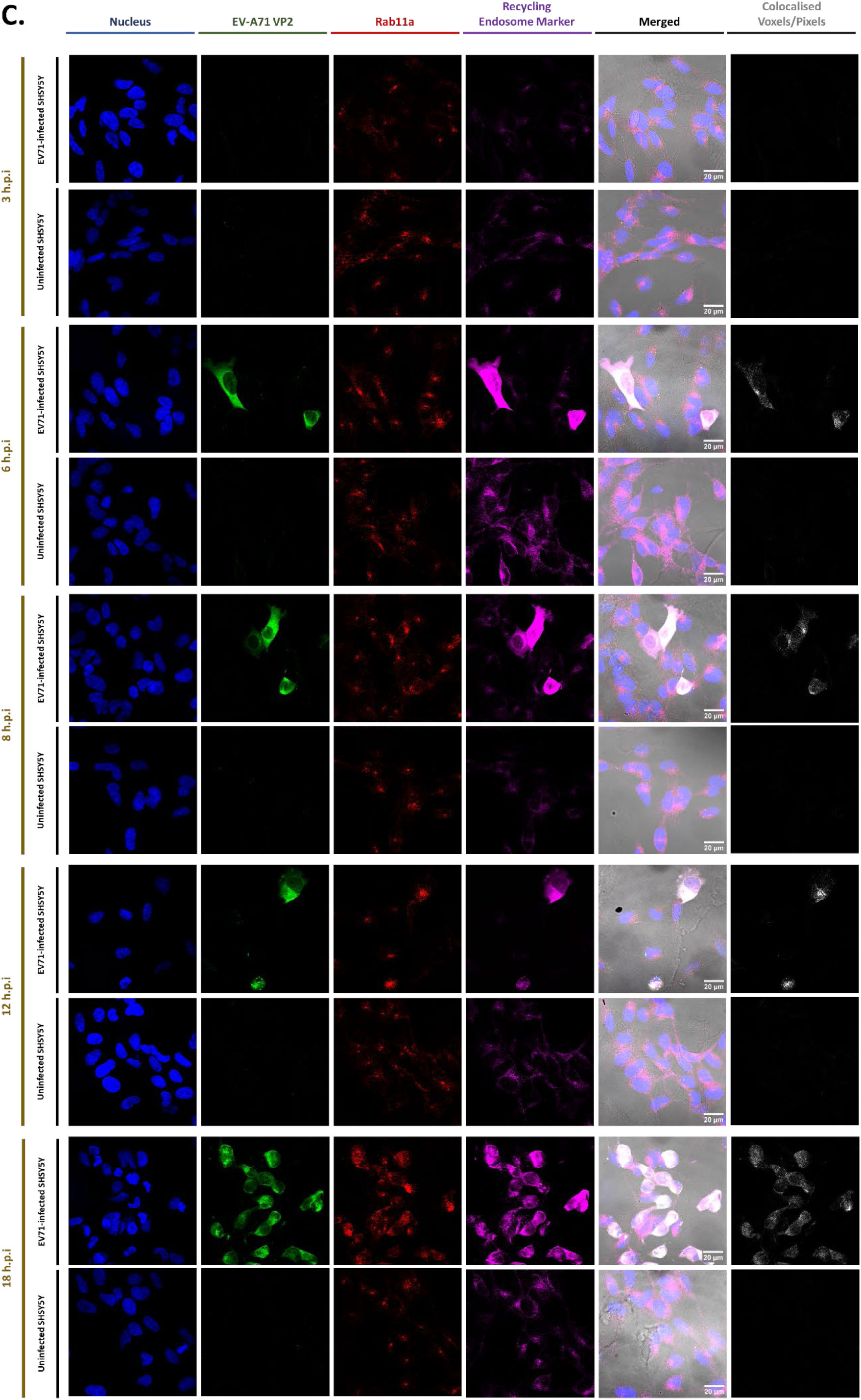
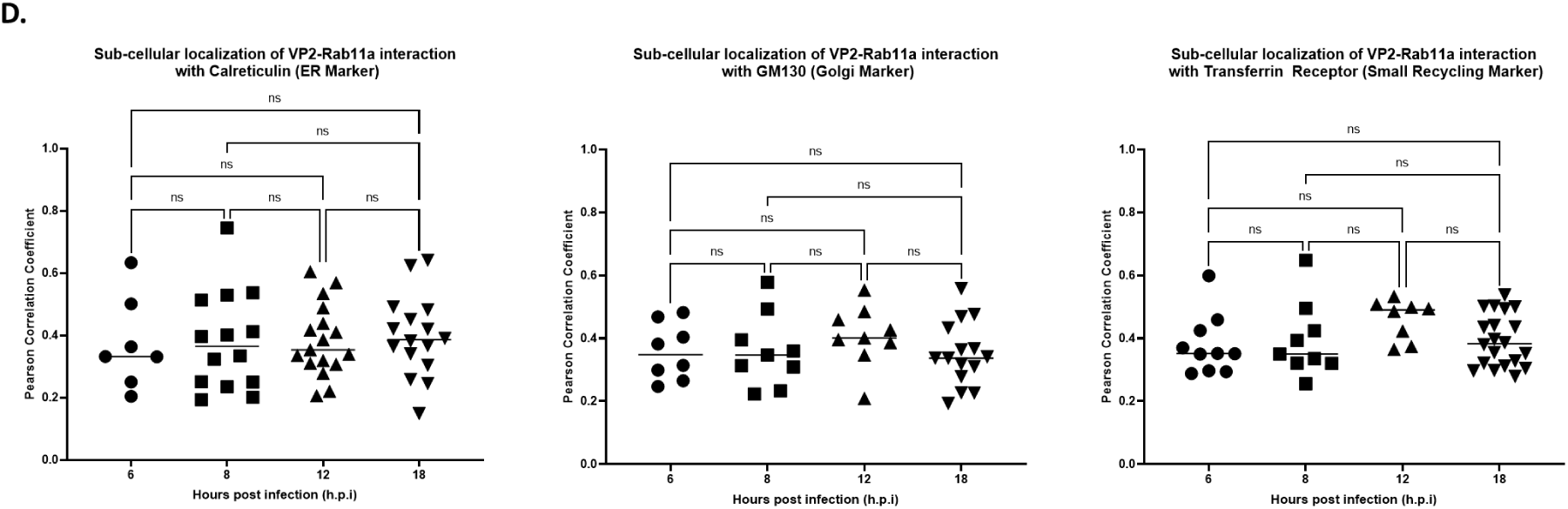
Colocalization between Rab11a and VP0/VP2 in various subcellular compartments over time. SH-SY5Y cells were infected with EV-A71 S41 (MOI 0.1). At the indicated time points post-infection, the cells were fixed and permeabilized, followed by staining with anti-Rab11a and anti-VP2 primary antibodies, and labelled secondary antibodies. Cells were then further stained with antibodies specific to compartment markers calreticulin for endoplasmic reticulum **(A)**, GM130 for Golgi apparatus **(B)** and transferrin receptor (Tfr) for small recycling endosomes **(C)** prior to DAPI staining. Confocal images were captured under 100X objective and were analyzed using Fiji software to determine the voxels/pixels that represent three channels (green, red, magenta) co-localization. Briefly, a mask representing the co-localization of Rab11a and each viral component was delineated and subsequently overlayed with compartment marker signals to generate the ‘co-localized voxels’ images shown on the far right column. **(D)** Pearson correlation coefficient (PCC) values were computed using Fiji software. PCC value of 0 = no co-localization; 0.1 – 0.3 = weak co-localization; 0.3 – 0.5 = moderate co-localization; 0.5-1 = strong co-localization. Statistical analysis was performed using Kruskal-Wallis test with Dunn correction against siNTC treatment (*p<0.05, **p<0.01, ***p<0.001, ****p<0.0001).

Altogether, this multiplex confocal imaging approach supported that during EV-A71 infection Rab11a interacts with all the viral components tested, as early as 6 h.p.i and across three major subcellular compartments. However, extensive remodelling of the ER and GA membranes during EV-A71 infection has been reported to form the replication organelles (RO), where major events of viral replication as well as viral morphogenesis occur [34]. It is hence possible that the observed Rab11a interactions with viral components may occur at the RO, and not at the ER and GA *per se*. To test this hypothesis, we analysed the co-localization between the three compartment markers during the course of infection. Cells were stained with a VP2 antibody to differentiate infected from uninfected cells in the culture. The images obtained indicated that at early time points during infection (before 6 h.p.i.) and in uninfected cells, minimal co-localization was observed between calreticulin, GM130 and Transferrin receptor (Fig. S5A). In contrast, from 6.h.p.i onwards, once VP0/VP2 can be readily detected, clear colocalization of the three markers was detected in the infected cells (Fig. S5A). PCC analysis confirmed significantly increased colocalization of the three markers in infected cells compared to uninfected cells (Fig. S5B). This finding therefore supported the extensive intracellular membrane remodelling during EV-A71 infection, where ER and GA merge to facilitate and support formation of replication organelles (RO). Consequently, it appears that interactions between Rab11a and the various viral components are likely to take place at the RO rather than in respective subcellular compartments.

### Role of Rab11 during EV-A71 infection is independent of its GTPase activity

Like all members of the Rab family, Rab11a/b contains a small GTPase domain that acts like a switch to cycle Rab11 between active (GTP-bound) and inactive (GDP-bound) states. Depending on its state, Rab11 interacts with distinct downstream effectors, driving distinct biological functions. Dominant negative (S25N; GDP-bound) and constitutively active (Q70L; GTP-bound) Rab11a mutants have been previously reported to study movement and fusion of Rab11a-containing membrane/vesicles [24, 35, 36]. Hence, we sought of using these Rab11a mutants to assess the importance of Rab11a GTPase activity during EV-A71 infection.

SH-SY5Y were transfected with Rab11aWT, Rab11aS25N and Rab11aQ70L plasmid constructs and were FACS sorted (thanks to a GFP tag) thereby enriching the cultures in transfected cells between 79 to 89%. GFP^+^ cells were then treated with siRab11b#9 (to knockdown specifically Rab11b expression), and infected with EV-A71. Both viral titers and VP2 signal intensities were comparable to those obtained with siNTC-treated cells (Fig. 6A&B), thus indicating that EV-A71 is able to exploit Rab11a regardless of its GTPase activity status and implying that EV-A71 does not exploit the trafficking function of Rab11a to shuttle viral proteins to various subcellular locations, a function that requires Rab11a GTPase activity [17].

**Figure 6.**
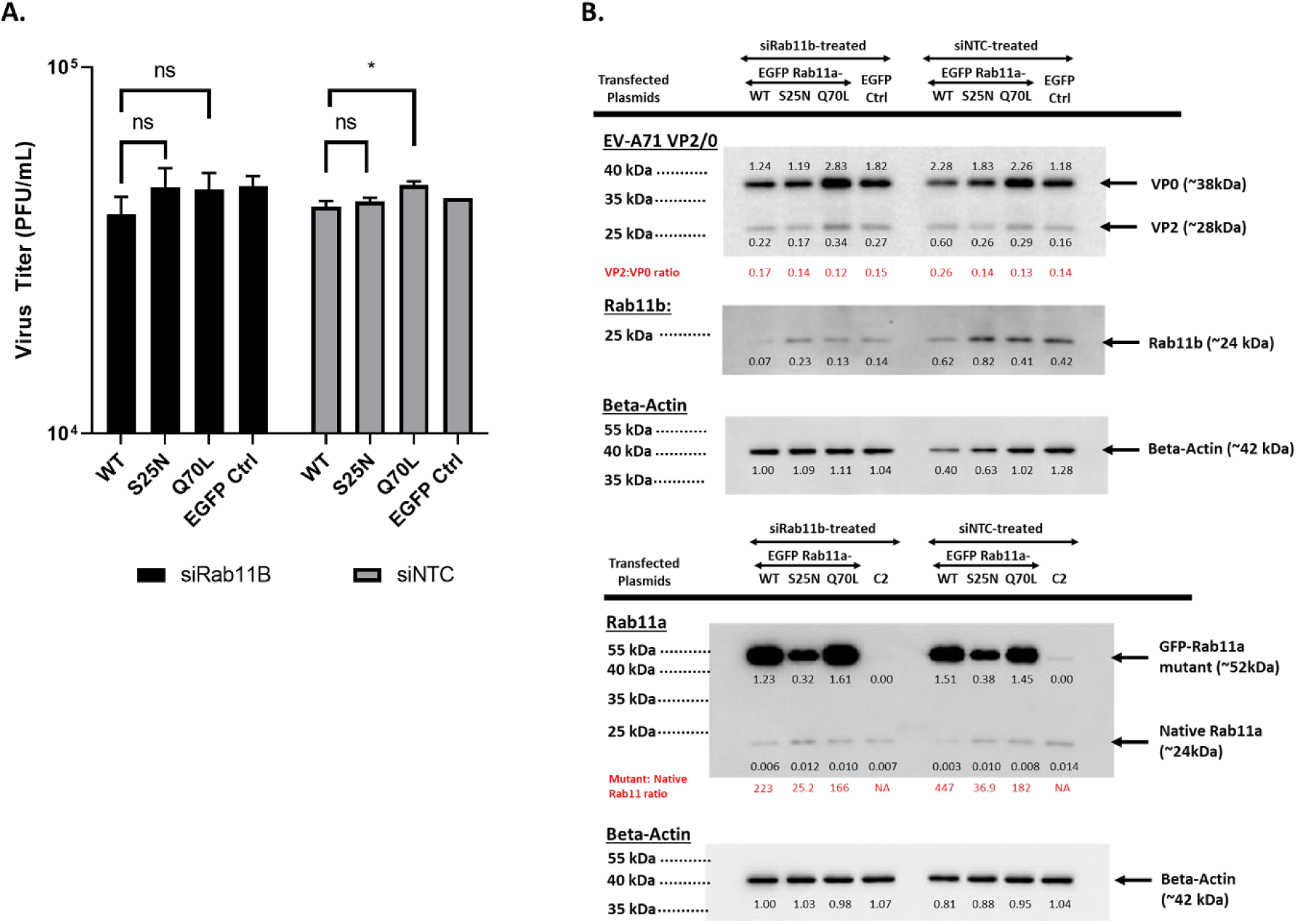
Effect of Rab11a DN and Rab11a CA overexpression on EV-A71 infection in siRab11b-knockdown SH-SY5Y cells. SH-SY5Y cells transfected with plasmids encoding for Rab11aWT, Rab11aS25N (dominant negative, DN), Rab11aQ70L (constitutively active, CA) and EGFP control, were FACS sorted (GFP+). The GFP+ enriched cell suspensions were then transfected with siRab11b #9 before infection with EV-A71 S41 (MOI 0.1). At 24 h.p.i, the culture supernatants and cell lysates were harvested. **(A)** Viral titers in culture supernatants were determined by plaque assay. Statistical analysis was performed using Mann-Whitney test against Rab11aWT (*p<0.05, **p<0.01, ***p<0.001, ****p<0.0001). **(B)** The cell lysates were analysed by Western blot using anti-Rab11a or anti-VP2 primary antibody. Beta Actin was probed for normalization purpose.

Furthermore, confocal microscopy indicated comparable colocalization patterns and strengths between Rab11aWT, Rab11aS25N and Rab11aQ70L with viral components (VP1, VP2, 3C, 3D, and dsRNA) (Fig. S6), further supporting that these interactions are GTPase activity-independent.

### EV-A71 re-directs Rab11a interactions with chaperones

To further characterize the role of Rab11a/b during EV-A71 infection, mass spectrometry of Rab11a pulldown (Fig. S7) from infected and uninfected samples was performed to identify Rab11a interacting partners. A total of 69 proteins were identified from the infected pulldown samples, far lesser than the 202 hits identified from the uninfected samples, with infected and uninfected samples sharing 50 common candidates (Fig. 7A; Table S4). Importantly, 19 of the 69 proteins found in infected samples were unique and not found in the uninfected pulldowns. These observations thus indicated that interactions between Rab11a and its host partners are significantly altered during EV-A71 infection. PantherDB classification revealed that the infected pulldown sample was enriched in cytoskeleton proteins, whereas the number of transporter proteins and proteins involved in membrane trafficking was greatly reduced compared to uninfected sample (Fig. 7B). HSPA8, HSPA2, CCT8 and CCT3 were identified as top hits that were greatly enriched in the infected pulldown sample (Table S5). These proteins fall in the category of chaperones and chaperonins, further supporting the scaffolding role of Rab11a during EV-A71 infection. HSPA8 has been previously suggested to participate in EV-A71 assembly/maturation, but with no known association with Rab11 in such process [37]. HSPA2 is a paralog of HSPA8 with 85% amino acid sequence identity. Since many HSPAs have overlapping functions [38], it is likely that HSPA8 and HSPA2 play a redundant role during EV-A71 infection.

**Figure 7.**
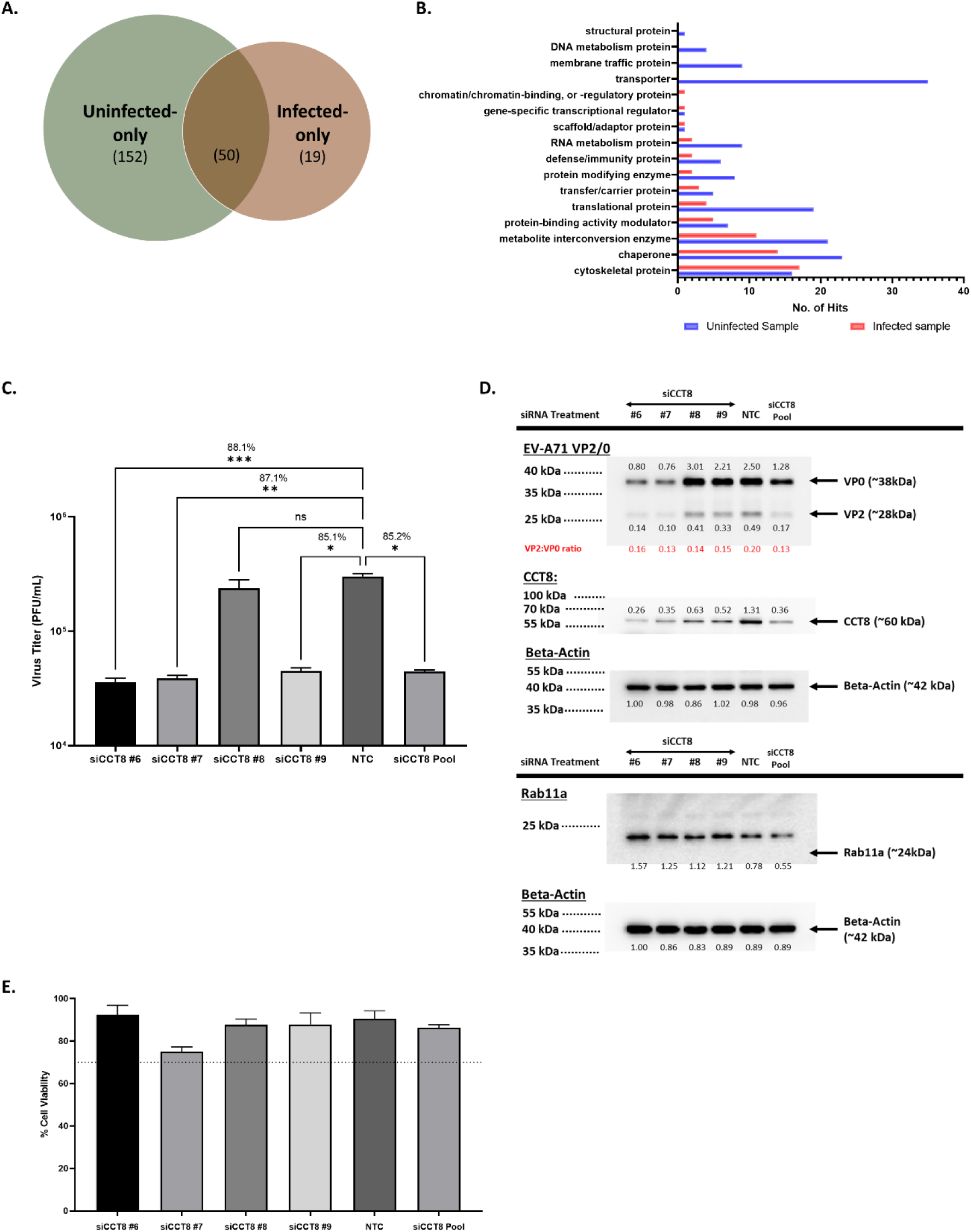
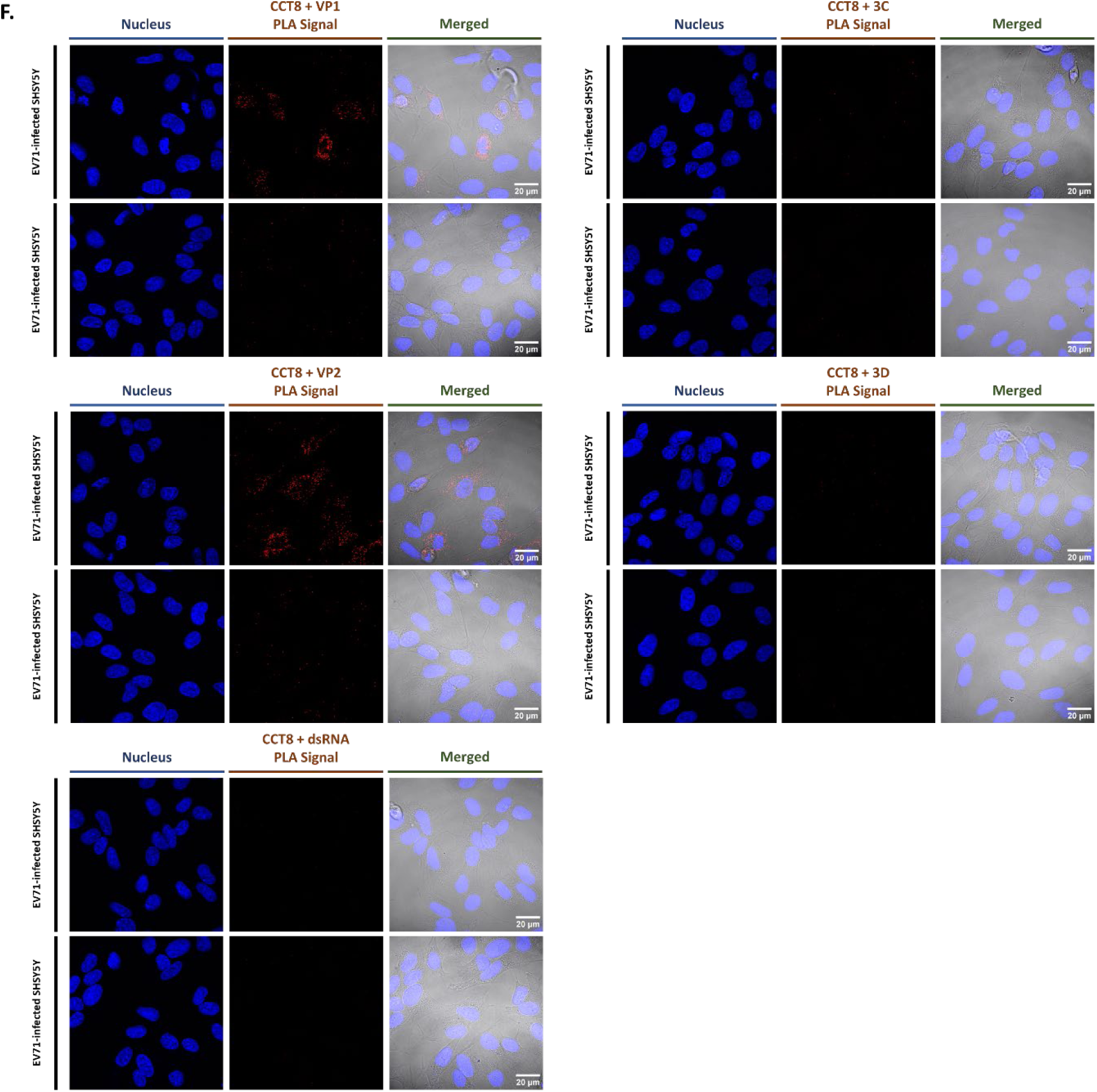
Rab11a interacting partners during EV-A71 infection. SH-SY5Y cells were infected with EV-A71 S41 (MOI 0.1). At 24 h.p.i, cell lysates were pulled down with anti-Rab11a antibody. The proteins were identified by mass spectrometry. **(A)** Venn diagram. **(B)** PantherDB classification of proteins only found or enriched in pulldowns from infected or uninfected samples. **(C,D)** SH-SY5Y cells were transfected with deconvoluted siRNA targeting CCT8 or siNTC. At 48h.p.t, cells were subjected to infection with EV-A71 S41 (MOI 0.1). At 24 h.p.i, the culture supernatants and cells were harvested. Virus titers in the cultured supernatants were determined by plaque assay (C). The percentage of viral titer reduction compared to NTC was indicated above the asterisk. Statistical analysis was performed using Kruskal-Wallis test against siNTC treatment (*p<0.05, **p<0.01, ***p<0.001, ****p<0.0001). Cell lysates were subjected to Western blot analysis using anti-VP2, anti-Rab11a, anti-CCT8 and anti B-actin antibodies (D). Band intensities were normalised to Beta-actin and were expressed relative to NTC. **(E)** AlamarBlue assay was performed in uninfected cells to assess cytotoxicity of siRNA treatments at 48 hours post-transfection. The readout for each treatment was relative to non siRNA-treated cells to determine the percentage of cell viability. Percentage above 70% (indicated by the dotted line) was considered non-cytotoxic. **(F)** Co-localization and interactions of CCT8 with viral components were assessed by proximity ligation assay (PLA). SH-SY5Y cells were infected with EV-A71 S41 (MOI 0.1). At 24 h.p.i., the cells were fixed and permeabilized before staining with anti-CCT8, paired with anti-VP2, anti-VP1, anti-3D, anti-3C or anti-dsRNA antibodies. The cells were further stained with secondary antibodies conjugated with DNA probes. Ligation and polymerase chain reaction were then carried out for signal amplification. Nuclei were stained with DAPI. Images were captured at 100X magnification under Olympus FV3000 confocal microscope.

On the other hand, involvement of chaperonins such as CCTs during EV-A71 infection has never been reported. siCCT8 KD led to decreased viral titers by approx. 1 log (Fig. 7C&D), thus supporting a pro-viral role for this chaperone. In addition, lower VP2:VP0 ratio was observed in siCCT8 KD samples, supporting a role for this chaperone in virus assembly/maturation (Fig. 7D). Interactions between CCT8 and various EV-A71 viral components were also assessed using PLA assay. Interestingly, positive PLA signal was detected when combining CCT8 with structural protein VP1 or VP2, but not with non-structural components 3C, 3D and dsRNA (Fig. 7F). This finding suggested the involvement of CCT8 in folding EV-A71 structural proteins to facilitate the assembly of provirion particles and/or induce conformational changes in the provirion essential for VP0 cleavage.

Together, the results indicated that during EV-A71 infection, Rab11a interactions with host factors were extensively redirected towards chaperones and cytoskeleton proteins, and away from proteins involved in transport and membrane trafficking. This observation further supported that Rab11a plays a minimal role in trafficking activities during infection but rather acts as an adapter or scaffold protein.

## DISCUSSION

Previous work has shown that host factors and pathways involved in membrane trafficking are exploited by enteroviruses to support various stages of their infection cycle. For example, Rab34 and Rab17 have been reported to contribute to coxsackievirus B entry step [39]. Mannose 6-phosphate receptors (MPRs), which are transmembrane glycoproteins that target enzymes to lysosomes, have been shown to be involved in EV-A71 uncoating [40]. SNARE proteins that drive membrane fusion and cargo exchange [41], were found to participate in EV-D68 genome replication as well as in virus exit [42]. SNARE SNAP29 has also been established as a pro-viral factor during EV-A71 infection [43]. To further our understanding on how EV-A71 harnesses membrane-associated pathways and factors, we screened a siRNA library comprising of 112 genes that are associated with membrane trafficking events. Rab11a was identified as a *bona fide* pro-viral factor in murine and human cell lines of neuronal and muscle origin. Our data strongly supported that Rab11a and Rab11b were used interchangeably by the virus. In addition, the role of these Rab11 proteins during infection was found to be conserved across EV-A71 sub-genogroups and CVA16, another major causative agent of HFMD.

A previous study reported the screening of a similar siRNA library in human intestinal Caco-2 cells during EV-A71 infection [44]. However, neither Rab11a nor Rab11b were identified as pro-viral factors with no significant reduction in viral titers in cells treated with siRab11a and siRab11b respectively [44]. The discrepancy between this earlier study and our work is likely attributable to the difference in cell line that was employed for the screening. Indeed, we showed that in human RD and SH-SY5Y cell lines, simultaneous siRNA KD of both Rab11a and Rab11b was necessary to observe viral titer reduction, due to functional redundancy between both proteins. This was not true in murine NSC34 cells where siRab11a KD alone was sufficient to observe viral titer reduction, although combined KD of Rab11a and Rab11b led to greater viral titer reduction. It is therefore very likely that combined KD of both Rab11 isoforms in Caco-2 cells would be required to observe significant impact on the viral titers. The different outcomes following Rab11a KD in various cell lines could stem from the differential relative expression of both isoforms in those cells.

Other RNA viruses including influenza and Ebola have been reported to exploit Rab11 proteins during their infection cycle [19, 21, 24]. Briefly, influenza virus was reported to exploit Rab11 to transport its ribonucleoproteins towards the plasma membrane where assembly and budding of newly formed viral particles occur [19, 24]. Similarly, during Ebola infection, Rab11 was shown to be involved in the transport of viral matrix protein VP40 towards the plasma membrane, for the release of newly made virus particles [21]. Current understanding of a possible role for Rab11 during enterovirus infection involves its interaction with viral protein 3A, and host proteins PI4KB and ACBD3, which were shown to participate to the re-programming of cholesterol shuttling processes during Poliovirus and Coxsackievirus B3 infection [25-29]. The authors proposed that those events allowed increase the intracellular pool of free cholesterol required for extensive membrane remodeling as part of the formation of replication organelles (ROs), where major viral processes occur including viral RNA synthesis, viral protein translation, and virus assembly and maturation. Consistently, a Super-Resolution 3D-SIM imaging approach showed that cholesterol from plasma membranes was re-distributed to Rab11-containing recycling endosomes that were re-directed to ROs during CVB3 infection [25]. Therefore, the authors of these studies proposed a role for Rab11 in the transport of free cholesterol to ROs during enterovirus infection.

Our work instead did not support a role for Rab11 in trafficking activities and membrane movements during EV-A71 infection. Several lines of experimental evidence supported this conclusion. Firstly, over-expression of dominant negative or constitutively active Rab11a mutants, which lock the protein in GDP- and GTP-bound state respectively, did not affect viral replication, strongly suggesting that Rab11 GTPase activity is dispensable during EV-A71 infection, while it is an absolute requirement for Rab11 involvement in the movement of recycling endosomes [16, 17]. Secondly, our confocal multiplex imaging approach indicated that Rab11 proteins interacted with all the viral components simultaneously and as early as 6 h.p.i. with no evidence of time-dependent interactions, which would have been expected should Rab11 be involved in transporting viral components across various subcellular locations. Further analysis of subcellular compartment markers indicated that those interactions likely occurred at the ROs.

Furthermore, we showed that Rab11 was not involved in the virus entry step (receptor-mediated endocytosis and viral uncoating), nor in viral protein translation and RNA genome replication. In contrast, our data supported that Rab11 may be involved in the virus assembly and maturation process as evidenced by lower VP2:VP0 ratio in cells treated with siRab11. Together, these findings led us to propose that Rab11 may act as a scaffold or adapter protein at the ROs for optimal assembly and VP0 cleavage-mediated maturation of newly formed viral particles. The identification of chaperones as top interacting partners of Rab11a during EV-A71 infection supported this view.

One possible explanation for the discrepancy between our study and earlier work describing the role of Rab11 in cholesterol shuttling processes during enterovirus infection [25], could be that while Rab11 is found on cholesterol-loaded recycling endosomes, it actually does not contribute to the movement of those recycling endosomes towards the ROs. Instead, this activity may be performed by other host factors. Alternatively, it is also possible that poliovirus and CVB3 exploit Rab11 proteins according to a different mechanism compared to EV-A71.

While little is known about the assembly and maturation processes of EV-A71 virus, our work seems to implicate Rab11’s involvement. Our model proposes that through re-directing of recycling endosomes to ROs, Rab11 proteins recruit a variety of chaperone proteins that together participate in the assembly and folding of newly formed virions, enabling VP0 cleavage. A recent report has described the involvement of HSPAs at all stages of EV-A71 infection cycle, with HSPA8 and HSPA9 specifically involved in the virus maturation step [37]. Consistently, we identified HSPA8 as one of the top interacting partners of Rab11a during EV-A71 infection, along with HSPA1 and HSPA2. More interestingly and uniquely, we validated Rab11a-interacting chaperone CCT8 as a *bona fide* pro-viral factor during EV-A71 infection. CCT8 is a component of chaperone complex TRiC/CCT, which plays a critical role in the folding of cytoskeleton proteins such as tubulin and actin [45]. TRiC consists of 8 CCT subunits that bind to different targets and substrates, while monomeric CCT subunits are functionally active too [45-48]. CCTs were found to be exploited by a number of viruses including reovirus and HCV for folding and stabilizing their viral proteins structure [49, 50]. A previous transcriptomic and proteomic study reported that CCT8 gene expression and protein levels were reduced in EV-A71 infected RD cells and the authors speculated that such reduction could favor cytoskeleton disruption and re-organization to facilitate formation of ROs [51]. Here, we propose that CCT8 monomers and/or TRiC may either be involved in folding individual viral structural proteins or induces conformational changes in newly formed Ev-A71 provirions to facilitate VP0 cleavage.

In conclusion, this work contributes to further our fundamental knowledge and understanding of the dynamic and complex molecular interactions between EV-A71 and its mammalian host. Given the lack of effective antiviral treatment to fight this disease, the identification of novel host factors exploited by EV-A71 represents a potential avenue for developing host directed therapies.

## Acknowledgement

We would like to express our gratitude to Prof. Kawaoka for sharing Rab11 S25N and Q70L mutant constructs; Dr Esther Koh and Ms Lee Su Yin for their assistance and training in confocal microscopy at LSI and NUS Medicine Confocal Microscopy Unit respectively; A/P Paul Hutchinson and his team for their expertise and assistance in cell sorting at LSI; the Proteomics and Mass Spectrometry Services core facility at Nanyang Technological University for the LC-MS/MS service.

## Author contributions

QYN, ZQL, and VM performed the experiments; JJHC and VTKC provided reagents; SKS analysed data; QYN and SA designed the experiments, analysed the data, and wrote the manuscript.

